# Computational Analysis of Structure and Binding Energy for Cisplatin-Loaded Proteins

**DOI:** 10.64898/2026.01.15.699809

**Authors:** Søren C. Spina, Joseph S. Bailey, Blaise Kimmel

**Author notes:** To whom correspondence should be addressed **Corresponding Author**, Blaise R. Kimmel, Ph.D., 151 W. Woodruff Avenue, Columbus, OH 43210.

## Abstract

Platinum-based drugs, such as cisplatin, are first-line chemotherapy treatments for patients with cancer. However, the success of these drugs is balanced with severe off-target toxicities and high dosing requirements, prompting the development of selective nanocarriers for targeted drug delivery. This study uses a computationally guided approach to examine the role of amino acids in cisplatin binding within proteins as nanocarriers. Using density functional theory, we quantify the binding of cisplatin to platinum-coordinating amino acids. We then rationally engineer a model MSH6 protein carrier, and evaluate the ability of MSH6 to bind cisplatin via molecular docking simulations. Structure predictions of the engineered MSH6 show that inserting the cisplatin-binding site has a limited impact on the nearby protein architecture of MSH6. Finally, we confirm and reveal cisplatin’s mechanism of action with DNA binding, and compare the energetic potentials of DNA binding from protein-delivered cisplatin to systemically administered cisplatin. Future studies will use these results to experimentally validate the binding of cisplatin in model protein carriers, and inform the strategic design and experimental development of a protein nanocarrier to achieve targeted drug delivery in cancer.

**TOC Figure:** 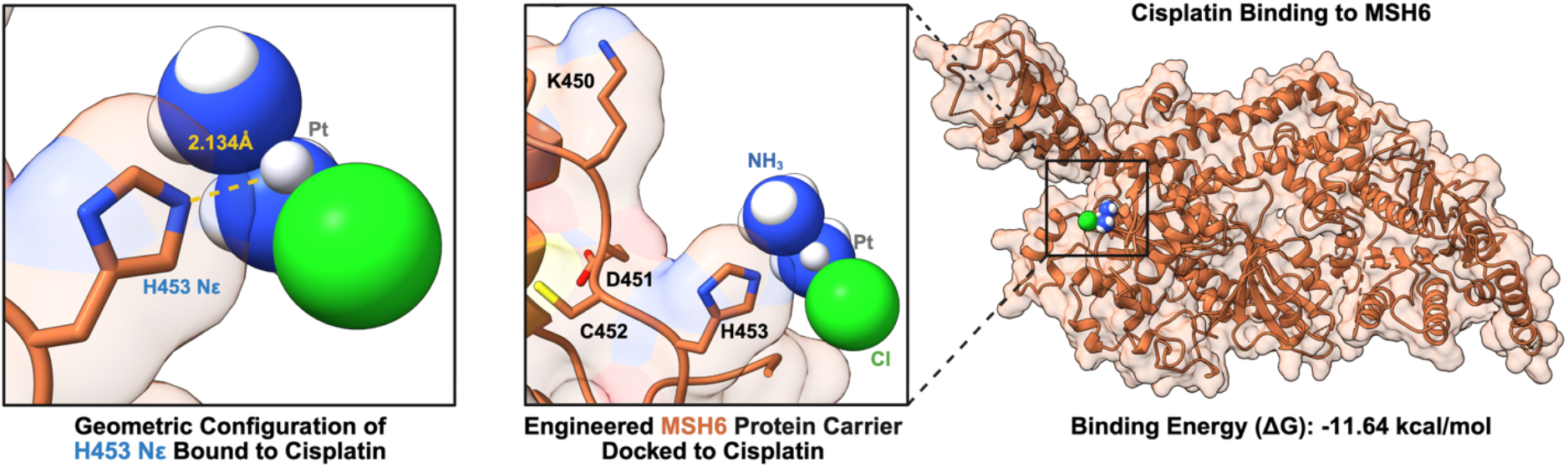

## Introduction

Cisplatin and platinum-based derivatives have been used to treat a wide range of cancer types, being a first-in-line treatment option for many sarcomas and carcinomas(1). When systemically administered to a patient, the amino groups typically enable cisplatin to enter cells via passive diffusion. In the cytoplasm, the amino groups of cisplatin undergo aquation, replacing the two bound chlorine atoms, forming dihydroxo moieties. In its water-soluble form, cisplatin exerts its chemotherapeutic action by crosslinking two purine bases on a DNA strand, causing irreparable damage that leads to cell death (**Figure 1**). Although cisplatin is highly effective at inducing apoptosis in tumor cells, systemic administration results in off-target cellular delivery, leading to DNA damage and toxicities in healthy tissues, especially the heart, liver, kidney, and hearing system(2). Many platinum-containing derivatives have been developed to exhibit potent anti-tumor effects via DNA damage, yet there remains a need for therapeutics that enhance the on-target delivery of these molecular agents.

**Figure 1.**
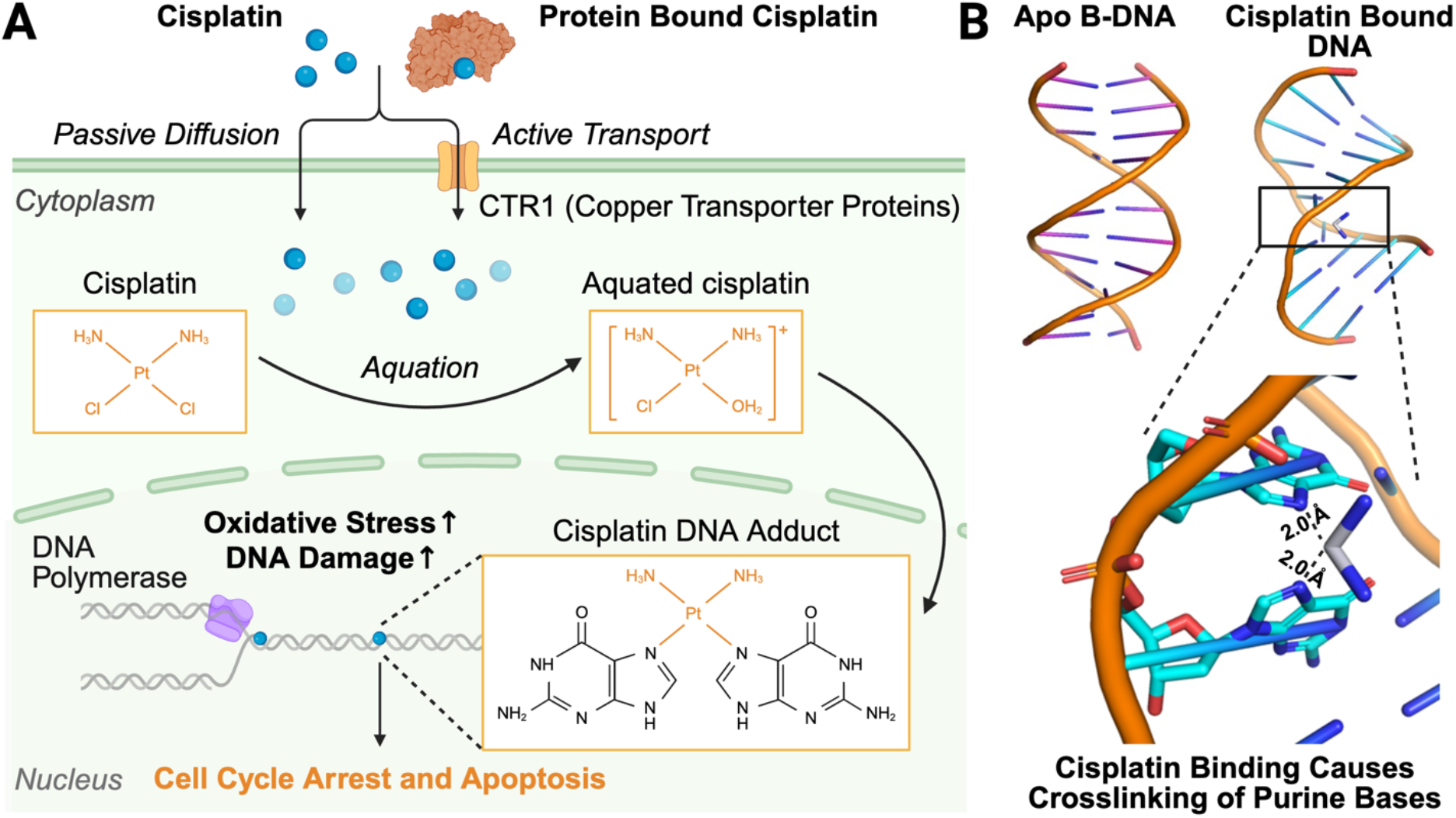
Cisplatin’s mechanism of action. (A) Cisplatin enters the cell via passive diffusion or active transport, undergoes aquation inside the cell, and forms a DNA adduct by (B) crosslinking purine bases, which can create bends (PDB: 1KSB) in straight DNA (PDB: 1AGH). Molecular visualization was performed with PyMOL.

Protein- and peptide-based therapeutics have recently gained popularity as potential vehicles for targeted delivery of molecular agents to treat cancer, including immune-activating drugs and chemotherapies, which have demonstrated significant improvements in on-target tumor delivery and reduced off-target toxicities in preclinical and clinical trials(3). By combining the selectivity of a protein therapy with the cytotoxicity of cisplatin, a selective and reliable cancer treatment could be developed for translation into the clinic. Supported by recent work in the field, apo-Human serum transferrin has been shown to improve tumor-specific delivery of cisplatin to human hepatocellular liver carcinoma cells *in vitro*, leading to apoptosis(4)’(5). Historically, human serum albumin (known for a long half-life in circulation) and the copper efflux transporter ATP7B have been observed binding to and transporting cisplatin to cells, where cisplatin is able to enter cells via passive or active diffusion, relying on cellular transport proteins (e.g., CTR1, copper transporter proteins) (**Figure 1A**)(6). While these examples demonstrate cisplatin’s ability to form a covalent adduct with proteins, there is a knowledge gap in predicting the binding site of cisplatin on candidate protein carriers and the residue to which it reacts, limiting the rational design of protein-drug conjugates for targeted cisplatin delivery.

This work addresses this gap by providing a computational investigation into how a biologic nanocarrier, at both the amino acid and protein level, plays a direct role in active cisplatin transport. We evaluate the binding interactions of key residues associated with cisplatin binding – histidine, lysine, methionine, arginine, cysteine, aspartate, and glutamate – to model cisplatin-protein binding via coordination of the platinum [II] ion. From these calculations, we rationally engineer a cisplatin-binding site into human MSH6, a model protein that binds to MSH2, thereby completing the formation of the MutSα complex(7). We use computational chemistry (via Density Functional Theory, DFT) calculations to study cisplatin binding and crosslinking purine nucleobases to understand how cisplatin then interacts with DNA, along with the role of aquation. Collectively, our study simulates and models the interactions between cisplatin and amino acids within a model protein, positioning cisplatin delivery as a therapeutic strategy via rational protein design.

## Computational Methods

### Cisplatin-Amino Acid Binding Calculations Using DFT

Cisplatin, monosubstituted aqua cisplatin, disubstituted aqua cisplatin, chloride, water, and amino acids were modeled in Avogadro 2, where an initial geometry optimization was completed using Babel (8). A full geometry optimization and frequency calculation was then performed in ORCA (9,10) using the close-shell restricted hybrid density functional B3LYP method with the SMD solvation model (11). For organic atoms, the 6-31+G(d,p) basis set was used. For the platinum atom, the def2-TZVP basis set with def2-ECP was used. A fine grid was used for numerical integration. Each multiplicity was set to 1, and the SCF was set as tight. The thermodynamics of cisplatin, monosubstituted aqua cisplatin, and disubstituted aqua cisplatin binding amino acids were calculated and recorded with the optimized geometry being modeled in Avagadro 2. The following equation was used to determine the total electronic energy (ΔE), enthalpy (H), and total free energy (ΔG).

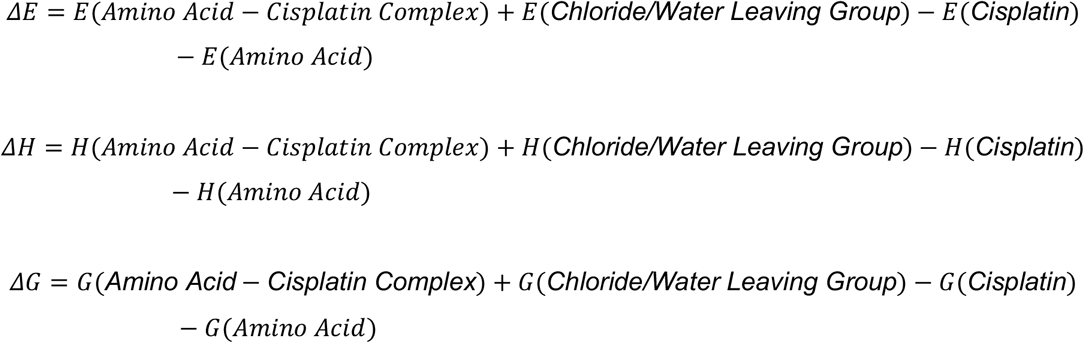

### *In Silico* Mutagenesis, Molecular Docking, and Structural Prediction of Engineered MSH6

We used MIB2: Metal Ion-Binding site prediction and modeling server to dock Pt^2+^ to the MSH6 crystal structure (PDB: 2O8B). Then, to begin input preparation for docking cisplatin to an engineered protein, the crystal structure of human MSH6 (PDB: 2O8B) was obtained from the Protein Data Bank and visualized in PyMOL version 3.0(12). Four residues (450-453) of the DNA-interaction site were mapped and mutated using the PyMOL mutagenesis tool to engineer a cisplatin binding site. This was repeated to create four engineered MSH6 variants (Mut: 1, Mut: 2, Mut: 3, and Mut: 4) , and the sequences of each are provided in the **Supplementary Information**. For each mutation, the rotamer of the lowest strain value was selected. Next, MetalDock was used to dock cisplatin against the MSH6 crystal structure along with the four engineered MSH6 structures. The pH was set to 7.5, with grid coordinates of the docking box being -46, 46, 36 and a box size of 19, 20, 20. Binding energies were recorded and used to determine the most favorable engineered MSH6 protein to proceed with. Boltz-2 and Chai-1, within the Tamarind Bio webserver, was used to predict the structure of engineered MSH6. With Chai-1 a multiple sequence alignment and the template crystal structure (PDB: 2O8B) were used for the structure prediction. With Boltz-2, a multiple sequence alignment, but no template crystal structures were used for structure prediction. Each predicted structure was aligned with the MSH6 crystal structure (PDB: 2O8B) using the ChimeraX Matchmaker tool.

### Cisplatin-Nucleobase Binding and Crosslinking Calculations Using DFT

Cisplatin, monosubstituted aqua cisplatin, disubstituted aqua cisplatin, chloride, water, and purine nucleobases were modeled in Avogadro 2, where an initial geometry optimization was completed using Babel (8). A full geometry optimization and frequency calculation was then performed in ORCA (9,10) using the close-shell restricted hybrid density functional B3LYP method with the SMD solvation model (11). For organic atoms, the 6-31+G(d,p) basis set was used. For the platinum atom, the def2-TZVP basis set with def2-ECP was used. A fine grid was used for numerical integration. Each multiplicity was set to 1, and the SCF was set as tight. The thermodynamics of cisplatin, monosubstituted aqua cisplatin, and disubstituted aqua cisplatin binding single and multiple nucleobases were calculated and recorded with the optimized geometry being modeled in Avagadro 2. The equation previously used to calculate the thermodynamics of amino acids binding cisplatin was used to calculate the thermodynamics of single purine nucleobases binding cisplatin. The following equation was used to determine the total electronic energy (ΔE), enthalpy (H) and total free energy (ΔG) of cisplatin crosslinking.

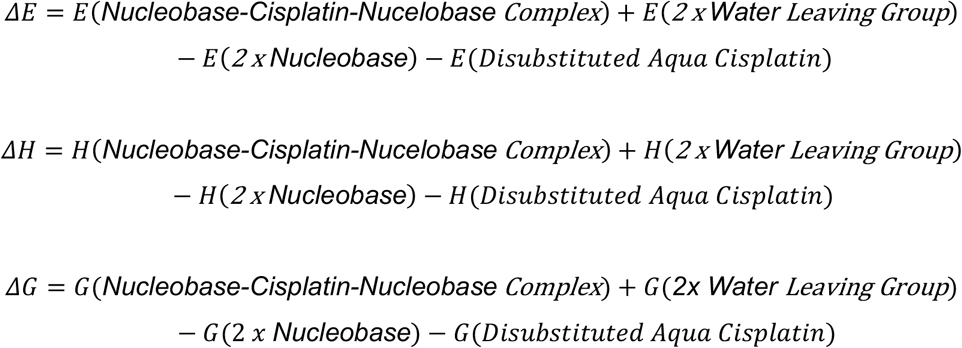

## Results

### Thermodynamics of Loading Cisplatin onto Amino Acids

We calculated and compared the binding abilities of individual amino acids to inform the rational design and generation of a cisplatin-binding pocket (**Figure 2**). Through observance of protein-cisplatin structures available on the protein data bank, histidine, lysine, methionine, cysteine, arginine, aspartate, and glutamate are the amino acids that directly coordinate the platinum atom of cisplatin. We used density functional theory with a B3LYP method within the ORCA program to calculate the electronic energy (E), enthalpy (H), and free energy (G) of amino acid-cisplatin complex formation. We performed these calculations with cisplatin, monosubstituted aqua cisplatin, and disubstituted aqua cisplatin to better inform the creation of a cisplatin binding site with tunable strength regarding aquation (**Table 1**).

**Table 1.** Calculated thermodynamics for amino acid binding disubstituted cisplatin. Inputs were modeled in Avogadro 2 and then geometrically optimized with a frequency calculation in ORCA via the restricted B3LYP method using a def2-TZVP basis set for platinum atoms and the 6-31+G(d,p) basis set for organic atoms.

| | $\Delta E_{\text{rxn}}(\text{kcal/mol})$ | $\Delta H_{\text{rxn}}(\text{kcal/mol})$ | $\Delta G_{\text{rxn}}(\text{kcal/mol})$ |
| --- | --- | --- | --- |
| **Amino Acid (AA) + Cisplatin → Cisplatin-Amino Acid + Chloride (Cl)** |  |  |  |
| Arginine | 15.14681 | 18.32766 | 24.31409 |
| Aspartate | -0.26795 | 1.009034 | 6.677323 |
| Cysteine | -16.4483 | -15.4486 | -10.5823 |
| Glutamate | -1.20369 | 1.046999 | 5.659064 |
| Histidine-Nε | 24.30487 | 25.3491 | 30.92176 |
| Histidine-Nδ | -1.20369 | 1.046999 | 5.659064 |
| Lysine | -8.68322 | -6.83702 | 0.169302 |
| Methionine | -0.91554 | 0.625062 | 7.08282 |
| **AA + Monosubstituted Aqua Cisplatin → Monosubstituted Aqua Cisplatin-AA + Cl** |  |  |  |
| Arginine | 19.60275 | 22.3594 | 30.22836 |
| Aspartate | -0.07907 | 0.599899 | 7.600389 |
| Cysteine | -14.776 | -13.2712 | -8.90498 |
| Glutamate | -0.2574 | 0.557856 | 6.688681 |
| Histidine-Nε | 19.82847 | 20.74087 | 27.92478 |
| Histidine-Nδ | 27.73998 | 29.22623 | 35.58786 |
| Lysine | -5.7156 | -3.15455 | 3.548814 |
| Methionine | 2.941762 | 4.631707 | 12.07905 |
| **AA + Disubstituted Aqua Cisplatin → Monosubstituted Aqua Cisplatin-AA + H<sub>2</sub>O** |  |  |  |
| Arginine | 4.208075 | 4.759028 | 9.843734 |
| Aspartate | -15.4737 | -17.0005 | -12.7842 |
| Cysteine | -30.1706 | -30.8716 | -29.2896 |
| Glutamate | -15.6521 | -17.0425 | -13.6959 |
| Histidine-Nε | 4.43379 | 3.140494 | 7.540148 |
| Histidine-Nδ | 12.3453 | 11.62586 | 15.20323 |
| Lysine | -21.1103 | -20.7549 | -16.8358 |
| Methionine | -12.4529 | -12.9687 | -8.30558 |

**Figure 2.**
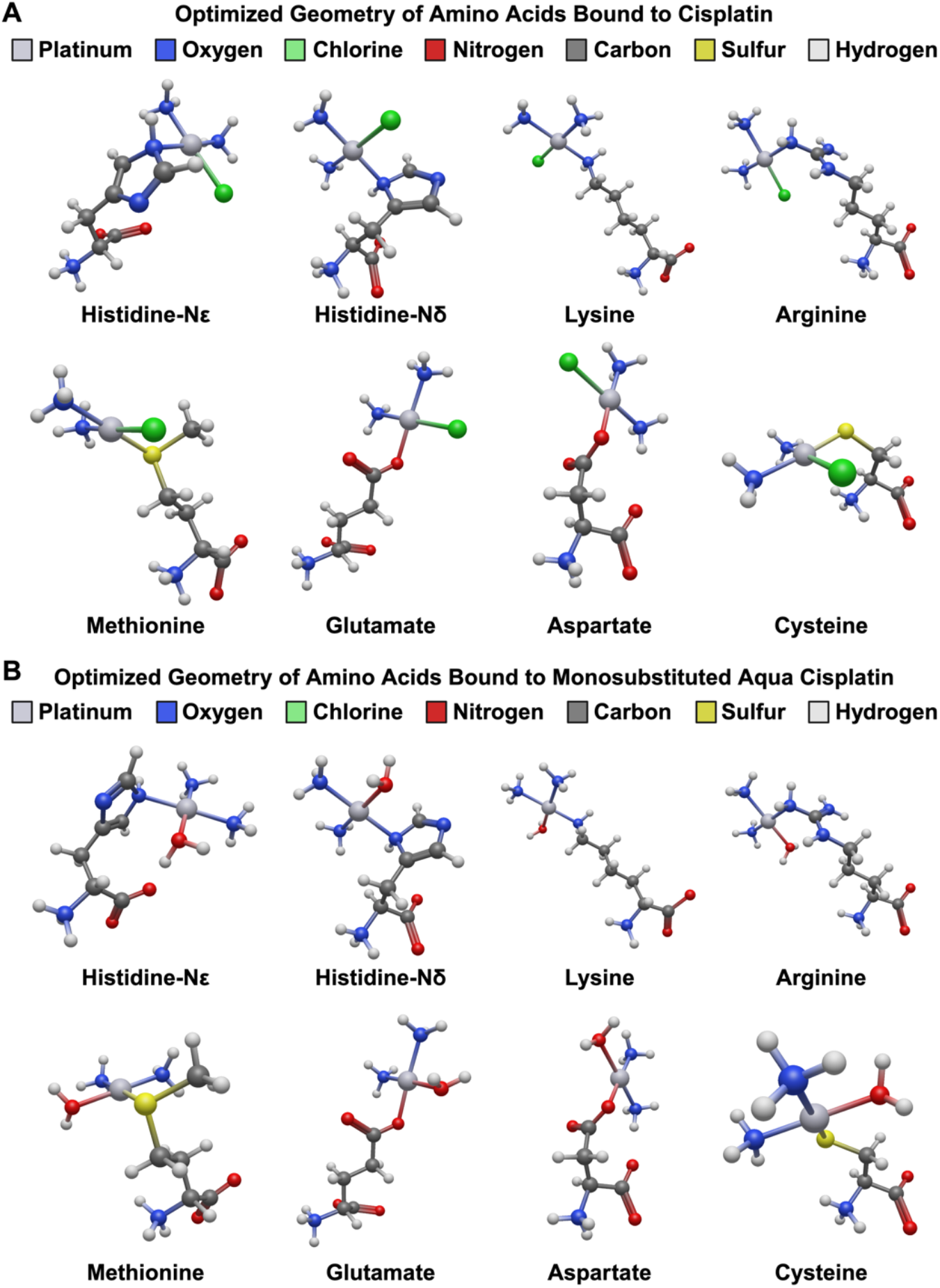
Geometrically optimized structures of cisplatin-binding amino acids. Optimized structures were modeled in Avogadro 2. Geometry optimization was performed with ORCA via the restricted B3LYP method using a def2-TZVP basis set for platinum atoms and the 6-31+G(d,p) basis set for organic atoms.

Histidine is the most prevalent amino acid that coordinates cisplatin in structures deposited in the Protein Data Bank. The platinum ion can coordinate to nitrogen at both the Nε and Nδ positions(13). We calculated that cisplatin binding, regardless of aquation status, was significantly more thermodynamically favorable for the ε-nitrogen than at the δ-nitrogen. This is likely due to steric interference of the protein chain on the residues. Histidine can bind cisplatin in a wide range of nitrogen-based coordination chemistries. With this in mind, and after analyzing the dynamic free energy results as aquation proceeds, we hypothesized that coordination to histidine is tunable depending on the nitrogen-binding site and the level of aquation. Further, this hypothesis is driven by the observation that histidine shares a functional group with the purine moiety in known cisplatin nucleobase binders, providing a potential analogue to examine thermodynamic differences in cisplatin coordination to the delta and epsilon nitrogen atoms of histidine.

Proteins have been shown to coordinate cisplatin with cysteine clusters, such as with the human copper chaperone, with four cysteine residues(14). After calculating the binding energy of cisplatin to cysteine, we found that it had the most thermodynamically favorable binding out of the amino acids tested. It bound disubstituted aqua cisplatin most strongly with an electronic energy and free energy that is ∼2x that of cisplatin and monosubstituted aqua cisplatin. The trend of cysteine favoring the disubstituted aqua cisplatin is consistent with previous computational studies(15). The other sulfur-containing amino acid, methionine, is also documented to coordinate cisplatin, for example, in the bovine pancreatic ribonuclease A adduct(16).

In previous studies, it has been shown that the thiol group of cysteine binds to cisplatin via charge transfer from the sulfur to the platinum atom in the shared covalent bond(15). While this interaction has been found to be strong for cysteine, similar reports from this work have suggested comparable ionic bonding between the sulfur atom in methionine and platinum. Further, some studies report that sulfur binding can induce ammonia release from the platinum [II] ion via the strong sulfur trans effect and trans influence, leaving cisplatin inactivated(17). Our work expanded on this, examining the chemical thermodynamics of bond coordination and electron density transfer between these sulfur atoms to platinum. Here, our DFT results suggest that cysteine coordination proceeds with a higher thermodynamic stability (ΔG_rxn,Cys_ = -10.5823 kcal/mol) compared to that of methionine coordination (ΔG_rxn,Met_ = 7.08282 kcal/mol) (**Table 1**).

Lysine has been shown to bind cisplatin in *Plasmodium falciparum* parasites, forming a reduced glutaredoxin 1 complex, as well as in thaumatin and human heavy chain ferritin(16)’(18). Lysine, Aspartate, and glutamate are similar amino acids that have been experimentally shown to coordinate cisplatin via an oxygen atom. These electrophilic atoms within the residue were expected to likely pair well due to the nucleophilic effects of the cisplatin binding to DNA. Aspartate and glutamate follow similar trends of binding, becoming thermodynamically favorable only when reacting with disubstituted aqua cisplatin.

### Studying Protein-Cisplatin Interactions with Molecular Docking

Over time, cisplatin has been adapted into numerous platinum-containing derivatives with significantly different structures. To investigate how platinum, as an ion, interacts with proteins, we used MIB2: Metal Ion-Binding site prediction and modeling server to investigate platinum binding from a whole-protein perspective(19). We docked Pt^2+^ against the MSH6 crystal structure and obtained binding scores for 22 different Pt^2+^ binding sites (**Table S1**). The individual acids consistently present in high-scoring binding sites are glutamate, methionine, and aspartate. The two highest-scoring sites involved coordination of methionine and glutamate (**Figure 3**) and the third highest-scoring site involved coordination between methionine and serine. Both calculations for amino acid-cisplatin binding and Pt^2+^ docking show interactions of glutamate and methionine, and to be very favorable.

**Figure 3.**
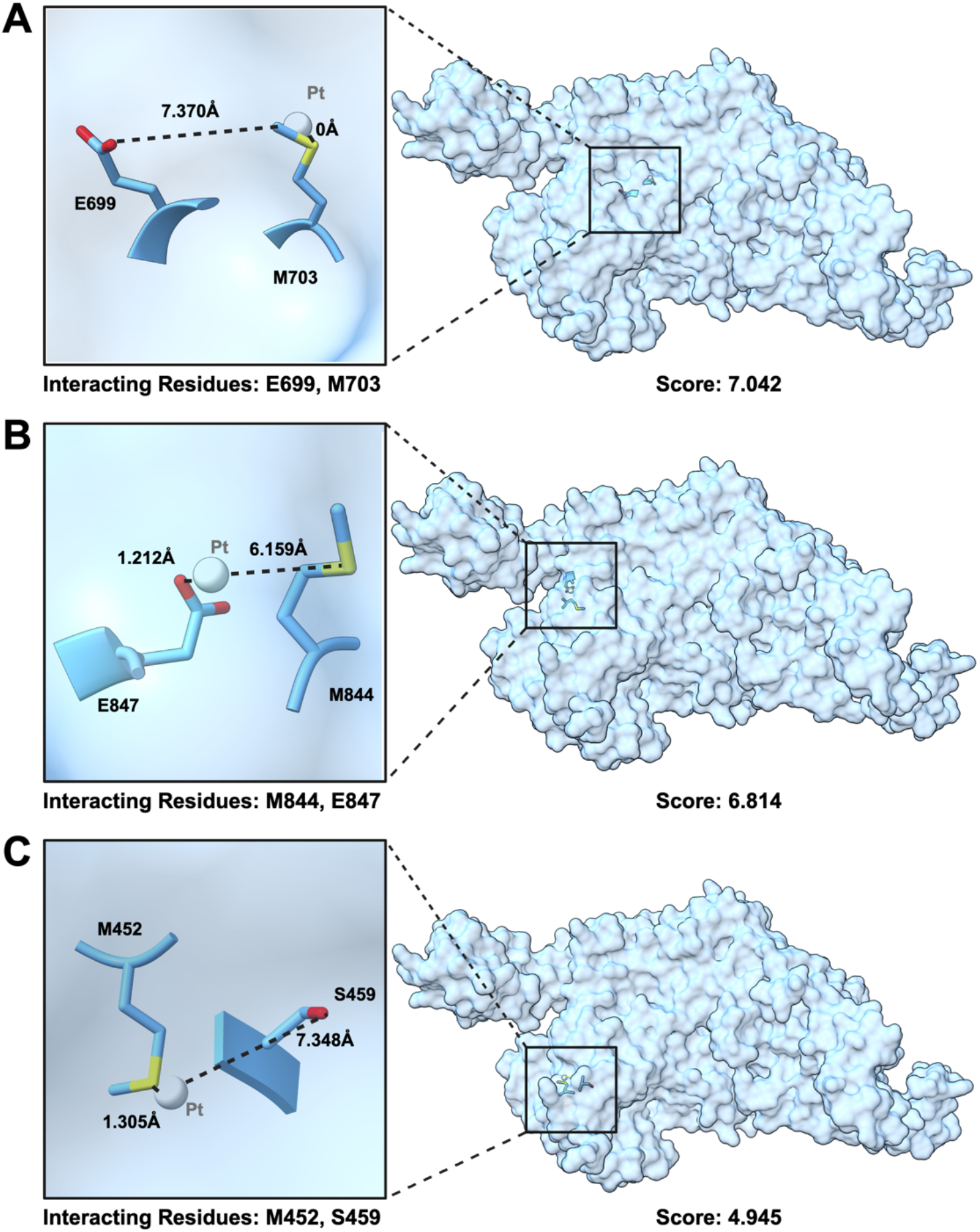
Docking of Pt^2+^ Against the Human MSH6 of the MutSα Complex. The MIB2 web server was used to dock Pt^2+^ to MSH6, computing coordination at (A) 844M and 847E, (B) 699E and 703, and (C) 452M and 459S. Atomic distances from Pt^2+^ to the residues are shown.

After observing how amino acids coordinate cisplatin and how protein binding sites coordinate platinum ions, we wanted to investigate how cisplatin binds at the protein level. We chose MSH6 as a model protein, from which the principles learned in this case study can be applied to engineer a cisplatin-binding site into any protein of interest that regularly interacts with DNA (**Figure 4A**). After observing experimental models of DNA binding by the MutSα complex, we chose four residues (450-453) with favorable proximity to purine-containing nucleobases for which we mutated to engineer a cisplatin-binding site (**Figure 4B**). The previous results on cisplatin-binding amino acids guided the rational design of this binding site, which comprises histidine, glutamate, lysine, and cysteine. Four engineered MSH6 versions were mutated so that each contained these four amino acids at different residue positions (**Table S2**). In this study, MSH6 serves as the model protein that is engineered to bind a chemotherapeutic payload; however, MSH6 itself does not exhibit tumor specificity.

**Figure 4.**
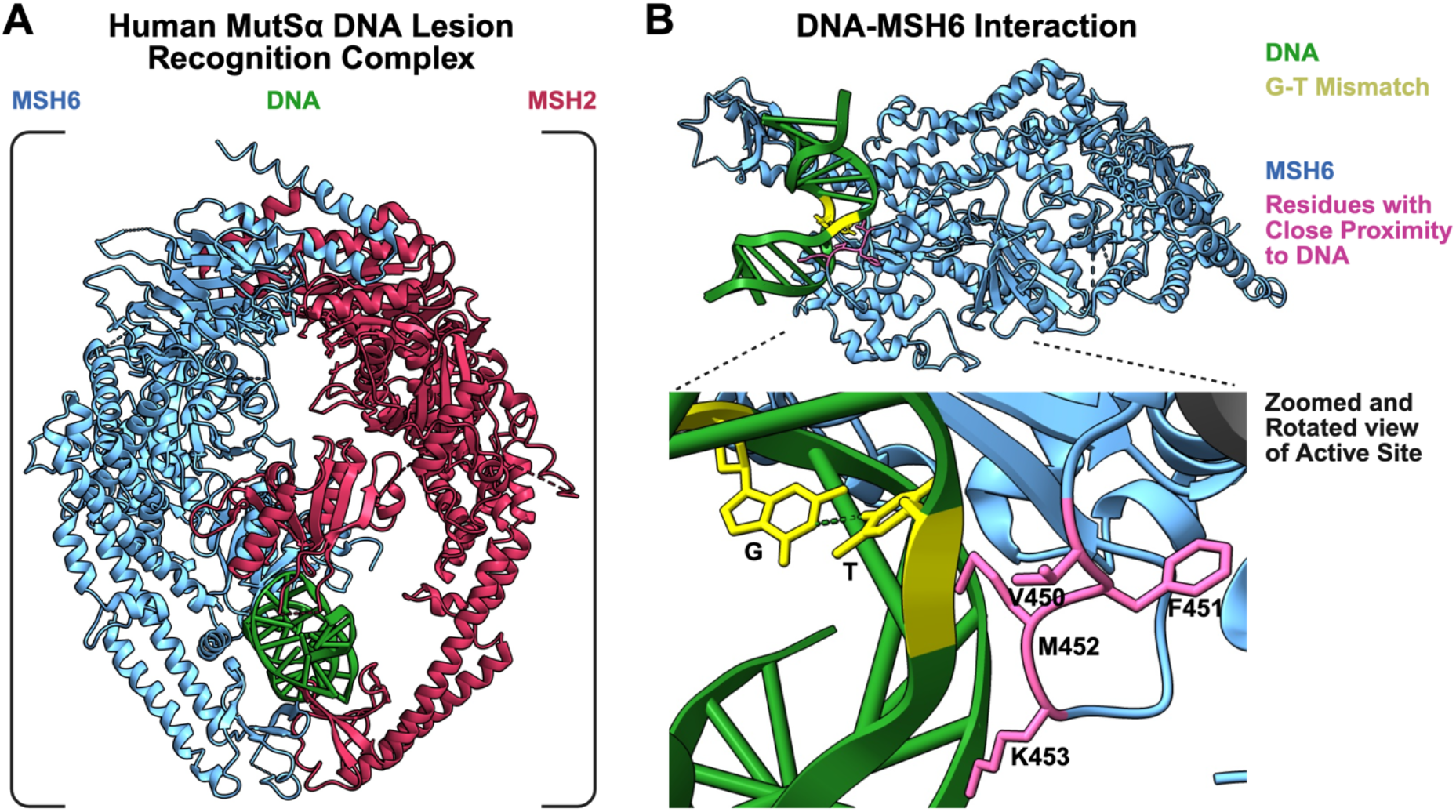
The Human MutSα Complex (MSH2/MSH6). (A) A DNA-lesion recognition complex (B) is sensitive to Guanine-Thymine base pair mismatches, PDB: (2O8B). Molecular visualization was performed with ChimeraX.

We used MetalDock to dock cisplatin to four rationally designed binding sites within an engineered MSH6. Each binding site differed in residue order but contained glutamate, histidine, lysine, and cysteine residues at positions 450-453. The most favorable engineered binding site, where cisplatin coordinated Histidine-Nε at the 453 position, was measured to have a binding energy of -11.64 kcal/mol (**Figure 5A**). It was unexpected that cysteine, lysine, and glutamate did not coordinate cisplatin during docking, despite these amino acids previously being calculated to individually bind cisplatin more favorably than histidine (**Table 1**). However, this result aligns with recent observations from experimental structures, as histidine is among the most common cisplatin binders in the Protein Data Bank(13). The other engineered sites found that interaction occurred between lysine and glutamate, yet not between cysteine. A potential reason for this is that cysteine is oriented toward the protein interior, where it is too small to accommodate cisplatin. Perhaps if the binding site were in a pocket rather than on an exterior loop, then cysteine would be the primary cisplatin binder. Another surprising observation was that cisplatin did not adopt a square-planar geometry within our most energetically favorable poses in the coordination and binding model. However, coordination could still be accounted for: the input cisplatin file coordinated two chlorine atoms, whereas the output file coordinated only one.

**Figure 5.**
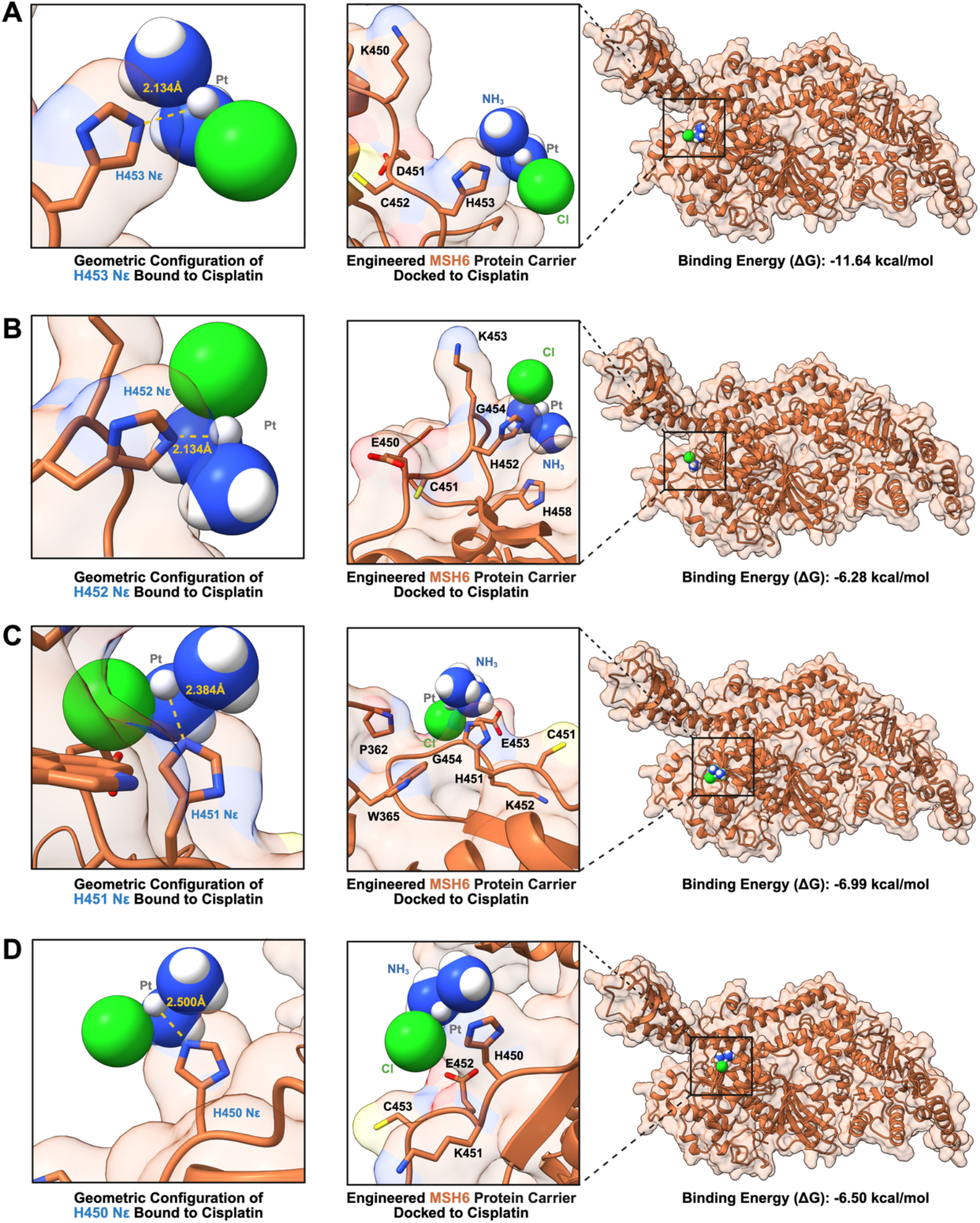
Molecular docking of cisplatin to engineered MSH6. Mutagenesis was performed on the MSH6 crystal structure (PDB: 2O8B), specifically residues V450, F451, M452, K453. MetalDock was used to dock cisplatin to engineered proteins with active sites made up of (A) K450, E451, C452, H453 (B) E450, C451, H452, K453 (C) C450, H451, K452, E453 (D) H450, K451, E452, C453. Each engineered protein carrier is visualized in a thermodynamically stable pose, as quantified by the negative binding energy (ΔG). Unmutated residues also were predicted to be involved with cisplatin binding in (B) G454, H458 and (C) P362, W365, G454.

### Predictive Analysis of the Structural Impact of Inserting the Cisplatin-Binding Site

We chose to insert the cisplatin-binding site into residues 450-453 of MSH6 because of its proximity to DNA within the MutSα complex. Additionally, this site is optimal because it is a loop rather than a part of the backbone secondary structure. Therefore, we hypothesized that mutagenesis of residues 450-453 would preserve the remaining MSH6 structure. To test this, we used Chai-1 with the MSH6 crystal structure template and Boltz-2 without a template to predict the structure of engineered MSH6 (**Figure 6**). The engineered MSH6 predictions and the MSH6 crystal structure were aligned in ChimeraX(20). The alignment score for the crystal structure against the Chai-1 predicted MSH6 was 4623.6, and the predicted Boltz-2 alignment was 4624.1 using the mutated ligand sequence outlined by residues 450-453.

**Figure 6.**
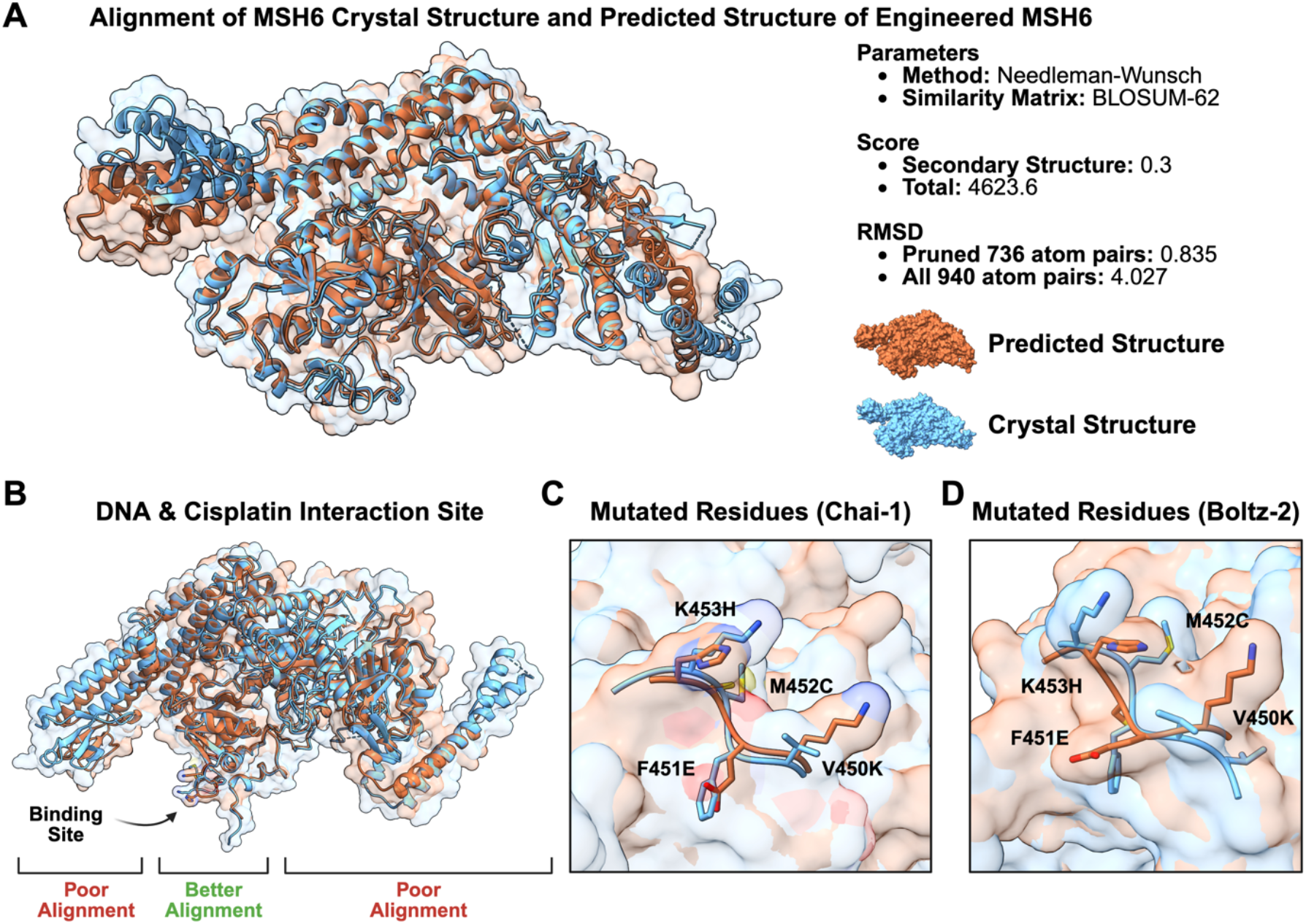
Alignment of MSH6 crystal structure to predicted structure of engineered MSH6. (A) Alignment of the MSH6 crystal structure to the structural prediction of engineered MSH6, using Chai-1. (B) Alignment of the Chai-1 predicted structure and the crystal structure, with better alignment at the cisplatin active site. (C) Mutated residues in alignment between Chai-1 predicted structure of MSH6 and MSH6 crystal structure. (D) Mutated residues in alignment between Boltz-2 predicted structure of MSH6 and MSH6 crystal structure.

A strong alignment is present at the active sites and progressively weakens farther from the active site. This observation was also evident during alignment of the unmutated MSH6 structure predictions with the MSH6 crystal structure, leading us to believe that the structural differences are more likely the result of imperfect structural prediction than of mutagenesis (**Figure S2**). Another potential reason is that gaps exist in the crystal structure that are not accounted for during structural prediction. The experimental properties of unmutated and each mutated MSH6 protein were predicted using Expasy ProtParam. The wild-type MSH6 and the engineered MSH6 with the strongest cisplatin-binding were predicted to be stable and to have long half-lives in yeast and mammalian cells. This provides initial evidence that these mutations do not significantly affect MSH6’s stability or compatibility with recombinant expression (**Table S5**).

### Confirm and Reveal the Mechanism of Action of Cisplatin with DNA

Cisplatin carries out its chemotherapeutic action by crosslinking two purine bases on a DNA strand, causing irreparable damage that leads to cell death. Many biochemical, biophysical, and structural studies indicate that crosslinking is the primary interaction of cisplatin with DNA(21,22). We used DFT calculations to investigate the optimized geometry and thermodynamics of cisplatin binding and crosslinking nucleobases across multiple cisplatin aquation states (**Figure 7**).

**Figure 7.**
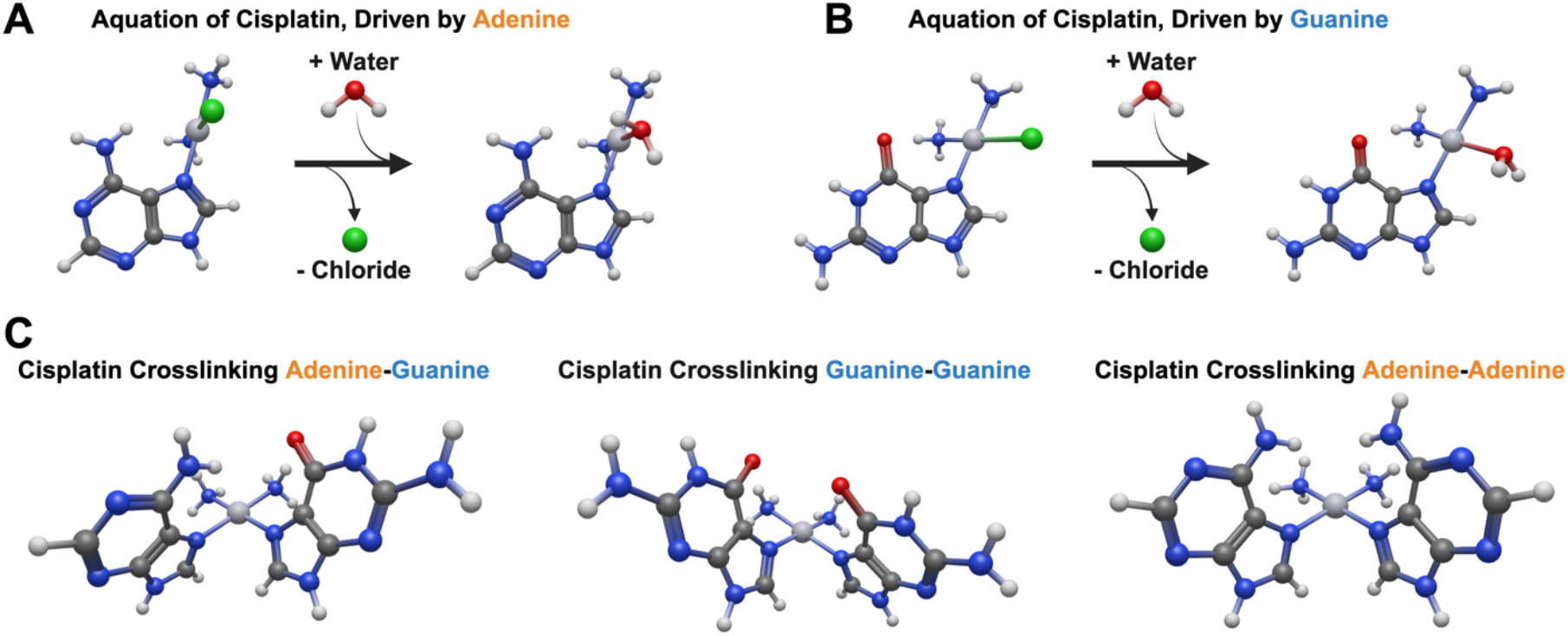
Geometrically Optimized Nucleic Acid Aquation and DNA Crosslinking. (A) Reactions associated with (A) adenine and (B) guanine driven aquation of cisplatin, yielding a monosubstituted cisplatin complex. (C) Cisplatin crosslinks two nucleic acids from the monosubstituted complex. Input and output structures were modeled in Avogadro 2. Geometry optimization was performed with ORCA via the restricted B3LYP method using a def2-TZVP basis set for platinum atoms and the 6-31+G(d,p) basis set for organic atoms.

Cisplatin undergoes aquation upon entering the cell, and it has been reported that the second aquation step occurs while bound to one nucleotide(23). Therefore, initially, monosubstituted cisplatin binds a nucleotide while kicking off a water ion, after which it undergoes a second aquation step before removing the chloride ion. The process of cisplatin crosslinking DNA has been thoroughly investigated both experimentally and computationally. By calculating the monosubstituted aqua-cisplatin interactions, we quantified the thermodynamics for each of the two possible reaction schemes. Here, we found that water was a favorable leaving group, whereas chloride was an unfavorable one, and that the binding of disubstituted cisplatin was slightly more thermodynamically favorable (**Figure 8**).

**Figure 8.**
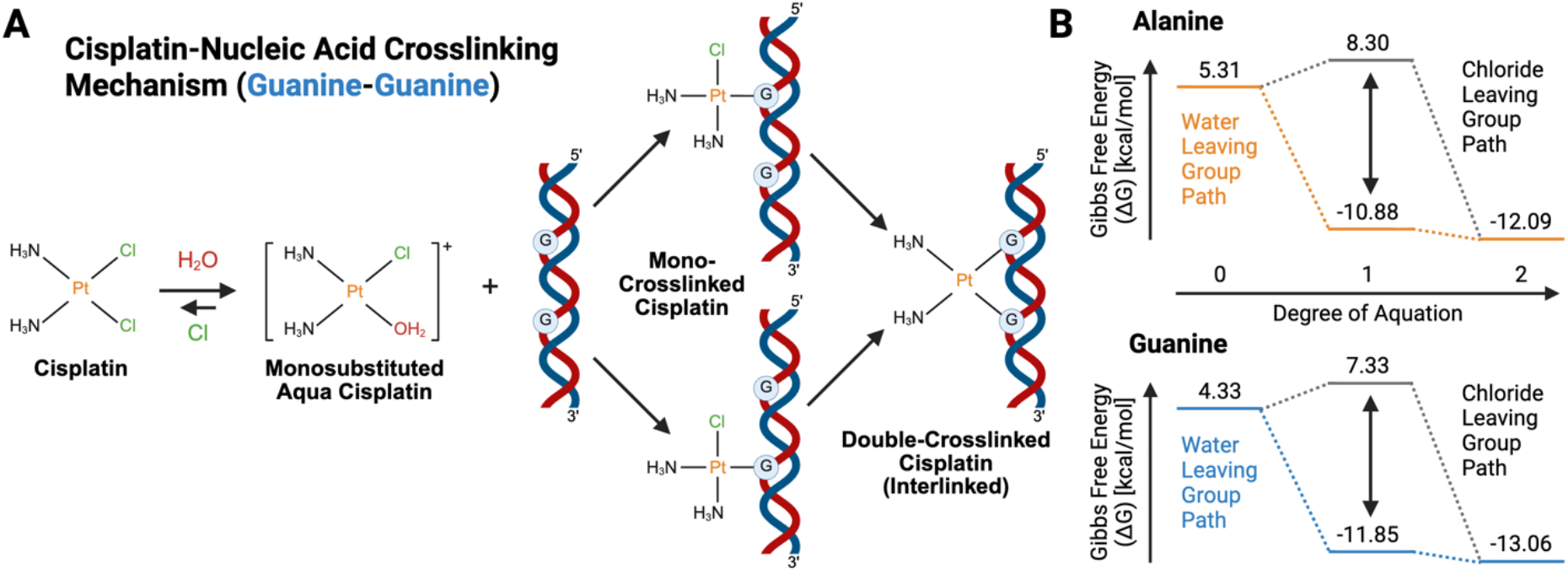
Mechanistic Insight into Cisplatin Aquation and Nucleobase Binding. (A) Reactions associated with cisplatin binding and crosslinking nucleobases. (B) When monosubstituted aqua cisplatin binds alanine and guanine, it is more thermodynamically favorable for water to serve as a leaving group than chloride.

Cisplatin crosslinking purine bases were modeled and thermodynamically investigated. Cisplatin crosslinking of two guanine bases was calculated to be the most thermodynamically favorable reaction, consistent with experimental findings(24). The thermodynamics of crosslinking adenine to guanine were found to be more favorable than crosslinking two adenine bases (**Figure 8**). In a model comparing the initial binding of cisplatin to either adenine or guanine, we found that guanine binding is significantly more favorable, again consistent with previously reported models and experiments(24). This trend was observed across all cisplatin aquation states (**Table 2**).

**Table 2.** Computationally derived thermodynamics for nucleobases binding cisplatin. Inputs were modeled in Avogadro 2 and then geometrically optimized with a frequency calculation in ORCA via the restricted B3LYP method using a def2-TZVP basis set for platinum atoms and the 6-31+G(d,p) basis set for organic atoms.

| | $\Delta E_{\text{rxn}}(\text{kcal/mol})$ | $\Delta H_{\text{rxn}}(\text{kcal/mol})$ | $\Delta G_{\text{rxn}}(\text{kcal/mol})$ |
| --- | --- | --- | --- |
| **Nucleobase + Cisplatin → Cisplatin-Nucleobase + Chloride (Cl)** |  |  |  |
| **Adenine** | -1.86031 | -0.99824 | 5.307973 |
| **Guanine** | -2.91233 | -1.51336 | 4.328181 |
| **Nucleobase + Monosubstituted Aqua Cisplatin → Monosubstituted Aqua Cisplatin-Nucleobase + Cl** |  |  |  |
| **Adenine** | 1.135478 | 2.054213 | 8.297991 |
| **Guanine** | -0.36244 | 0.37702 | 7.326431 |
| **Nucleobase + Monosubstituted Aqua Cisplatin → Monosubstituted Aqua Cisplatin-Nucleobase + H<sub>2</sub>O** |  |  |  |
| **Adenine** | -13.66 | -14.8674 | -10.8736 |
| **Guanine** | -14.712 | -15.3826 | -11.8534 |
| **Nucleobase + Disubstituted Aqua Cisplatin → Monosubstituted Aqua Cisplatin-Nucleobase + H<sub>2</sub>O** |  |  |  |
| **Adenine** | -14.2592 | -15.5462 | -12.0866 |
| **Guanine** | -15.7571 | -17.2234 | -13.0582 |
| **2 Nucleobases + Disubstituted Aqua Cisplatin → Nucleobase-Cisplatin-Nucleobase + 2 H<sub>2</sub>O** |  |  |  |
| **Adenine-Cisplatin-Adenine** | -27.8521 | -28.5445 | -21.205 |
| **Adenine-Cisplatin-Guanine** | -30.3336 | -31.382 | -24.361 |
| **Guanine-Cisplatin-Guanine** | -30.5517 | -31.7321 | -25.1664 |

## Conclusion

In this work, we evaluated the mechanism of action of cisplatin, from its coordination with a model protein carrier to its reaction with purine nucleobases, quantifying the energetics and thermodynamics of cisplatin binding. Our docking studies of engineered human MSH6 reveal preferential cisplatin binding to histidine, where the binding pocket contains cysteine, glutamate, and lysine residues as key neighbors of the cisplatin complex. These results are surprising compared to DFT calculations of amino acids, which showed a stable, preferential sulfur-based platinum coordination with cysteine. The cysteine coordination was the strongest among the amino acids, regardless of the cisplatin aquation state. This observation of high-affinity capture to cysteine is consistent with previous experimental reports that cysteine is a preferred amino acid for covalent adduct formation(24). A potential reason why our cysteine residue didn’t coordinate cisplatin is its orientation. Future work can repeat this protocol, but with amino acid residue positions swapped.

Further, we considered the potential for nitrogen-based coordination chemistries, hypothesizing that coordination to histidine is tunable depending on the nitrogen-binding site and the degree of aquation. Histidine shares a functional group with the purine group in known nucleobase binders of cisplatin, providing a potential analogue for understanding thermodynamic differences in cisplatin coordination to the delta and epsilon nitrogen atoms of histidine. Balancing stability during transport to the cell via proteins and release within the cell, histidine provides a novel strategy to modulate the strength of the protein-cisplatin interaction, thereby enhancing cellular uptake and increasing the rate of release from the protein within the cell. Once inside the cell and unbound from the protein carrier, we observed a strong thermodynamic drive for cisplatin to crosslink DNA, consistent with previous reports. Specifically, we found that for purine nucleobases (i.e., adenine and guanine), the disubstituted aquation complex of cisplatin is most thermodynamically favorable. Among the purine nucleobases, we observed that cisplatin preferentially binds to guanine rather than adenine, and the bridged guanine-cisplatin-guanine complex was the most favorable configuration, reinforcing the mechanistic endpoint that cisplatin aquation is thermodynamically most favorable for nucleic acid interactions.

## Discussion

These results provide a rational path from amino acid energetics to the design of active sites within protein candidates for chemotherapy delivery. Here, these active sites can be mutated to modulate cisplatin binding affinity by tuning the rate of aquation, altering structural accessibility within the carrier scaffold, and orienting cysteine or histidine residues (as sulfur vs nitrogen donors) at the site, thereby strengthening the protein-cisplatin interaction and enhancing cellular uptake. While high-affinity chemistries are sought after in the protein-drug conjugate field, a balancing act may be required for optimal targeted delivery of cisplatin, in which stable loading of the platinum coordinator should be balanced with kinetic trapping (i.e., premature sequestration) in off-target proteins. While cysteine is traditionally the active-site binder of cisplatin, forming a histidine-centered coordination of cisplatin offers a unique strategy to build carriers that enhance the circulation half-life and tumor delivery of cisplatin from systemic administration, while still allowing rapid aquation upon release from the protein carrier and DNA adduct formation.

Future studies can build off this work by creating a fusion construct between MSH6 and a delivery protein, such as an antibody or nanobody(25,26). Fusion proteins have recently shown success in localizing enzyme function in HER2^+^ breast cancer mouse models(27) and in selectively delivering STING-pathway-activating proteins(28)’(29). Further, in the design of protein carriers, it is important to recognize that the modeled protein, MSH6, may have limited transport properties after administration due to its size and solubility, even when fused to a nanobody. In that event, knowledge of cisplatin-binding amino acids could be applied to install a platinum-drug-binding site into a different or artificial protein scaffold. We expect that, from these results, rational protein design will play a key role in creating tunable cisplatin-binding sites within optimal protein carriers for low-toxicity, tumor-specific delivery (e.g., engineered nanobodies for high-affinity tumor delivery)(29–31) and in providing a strategy for integrating an expanded library of cisplatin derivatives.

## AUTHOR INFORMATION

## Author Contributions

Soren Spina: Wrote the original draft of the manuscript, generated all figures and graphics for the manuscript, edited, revised, and approved the final version of the manuscript. Joe Bailey: Supported data generation for the manuscript and figure preparation. Blaise Kimmel: Edited, revised, and approved the final version of the manuscript, and acquired funding to support the work.

### Funding Sources

We gratefully thank the Ohio State University Comprehensive Cancer Center (OSUCCC), OSUCCC Center for Cancer, and the Department of Chemical and Biomolecular Engineering at The Ohio State University for support of this work.

## Acknowledgements

This work was supported in part by The Ohio State University Center for Cancer Engineering-Curing Cancer Through Research in Engineering and Sciences. B.R.K. acknowledges financial support from the Prostate Cancer Foundation Young Investigator Award. All figures created with Biorender.com.

