## Supplemental Information for "Computational Analysis of Structure and Binding Energy for Cisplatin-Loaded Proteins"

*To whom correspondence should be addressed

### Supplementary Tables

**Table S1. Prediction Scoring of 23 different $Pt^{2+}$ binding sites.**

| No. | Binding Residues | Template | Score |
|---|---|---|---|
| 1 | 699E, 703M | 4lq6A0 | 7.042 |
| 2 | 844M, 847E | 3m7kA0 | 6.814 |
| 3 | 452M, 459S | 4lq6A0 | 4.945 |
| 4 | 934D, 936D, 937Y | 6mw5A0 | 4.625 |
| 5 | 665E, 667D | 6mw5A0 | 4.587 |
| 6 | 580S, 1202M | 5mu1A2 | 4.224 |
| 7 | 544E, 601L | 4pkeA0 | 4.007 |
| 8 | 703M, 826H | 4dpeA0 | 3.98 |
| 9 | 826H, 1042K | 6aqkA0 | 3.544 |
| 10 | 392D, 394S | 6ig4A0 | 3.503 |
| 11 | 380D, 381E | 6ig4A0 | 3.107 |
| 12 | 547E, 550S | 6ig4A0 | 3.027 |
| 13 | 562D, 564S | 6ig4A0 | 3.025 |
| 14 | 839D, 844M | 4dpeA0 | 2.893 |
| 15 | 380D, 382H, 383R | 6mw5A0 | 2.835 |
| 16 | 576D, 578H, 581R | 6mw5A0 | 2.814 |
| 17 | 826H, 830S | 6fhnA1 | 2.709 |
| 18 | 1250H, 1254E | 4pm3A0 | 2.64 |
| 19 | 1080L, 1119E | 1rnlA0 | 2.496 |
| 20 | 833K, 837H | 3w3sA0 | 2.477 |
| 21 | 761R, 1153M | 4ombC0 | 2.338 |
| 22 | 629D, 631S | 6ig4A0 | 2.3313 |

**Table S2: Sequences of MSH6 (Blue are sites of mutagenesis, yellow is mutagenesis).**

| | |
|---|---|
| MSH6<br>(Truncated<br>for<br>expression,<br>from PDB:<br>2O8B | PTVWYHETLEWLKEEKRRDEHRRRPDHPDFDASTLYVPEDFLNSCTPGMR<br>KWWQIKSQNFDLVICYKVGKFYELYHMDALIGVSELGL**VFMK**GNWAHSGFP<br>EIAFGRYSDSLVQKGYKVARVEQTETPEMMEARCRKMAHISKYDRVVRREI<br>CRIITKGTQTYSVLEGDPSENYSKYLLSLKEKEEDSSHTRAYGVCFVDTSLGK<br>FFIGQFSDDRHCSRFRTLVAHYPPVQVLFEKGNLSKETKTILKSSLSCSLQEG<br>LIPGSQFWDASKTLRTLLEEEYFREKLSDGIVMLPQVLKGMTSESDSIGLTPG<br>EKSELALSALGGCVFYLKKCLIDQELLSMANFEEYIPLDSDTVSKAYQRMVLD<br>AVTLNNLEIFLNGTNGSTEGTLLERVDTCHTPFGKRLLKQWLCAPLCNHYAIN<br>DRLDAIEDLMVVPDKISEVVELLKKLPDLERLLSKIHNVGSPLKSQNHPDSRAI<br>MYEETTYSKKKIIDFLSALEGFKVMCKIIGIMEEVADGFKSKILKQVISLQTKNP<br>EGRFPDLTVELNRWDTAFDHEKARKTGLITPKAGFSDYDQALADIRENEQSL<br>LEYLEKQRNRIGCRTIVYWGIGRNRYQLEIPENFTTRNLPEYELKSTKKGCKR<br>YWTKTIEKKLANLINAEERRDVSLKDCMRRLFYNFDKNYKDWQSAVECIAVL<br>DVLLCLANYSRGGDGPMCRPVILLPEDTPPFLELKGSRHPCGDDFIPNDILIG<br>CEEEEQKAYCVLVTGPNMGGKSTLMRQAGLLAVMAQMGCYVPAEVCRLTP<br>IDRVFTRLGSTFFVELSETASILMHATAHSLVLVDELGRGTATFDGTAIANAVV<br>KELAETIKCRTLFSTHYHSLVEDYSQNVAVRLGHMACMTFLYKFIKGACPKS<br>YGFNAARLANLPEEVIQKGHRKAREFEKMNQSLRLFRE |
| MSH6 Mut: 1<br><br>450K<br>451E<br>452C<br>453H | PTVWYHETLEWLKEEKRRDEHRRRPDHPDFDASTLYVPEDFLNSCTPGMR<br>KWWQIKSQNFDLVICYKVGKFYELYHMDALIGVSELGL**KECH**GNWAHSGFP<br>EIAFGRYSDSLVQKGYKVARVEQTETPEMMEARCRKMAHISKYDRVVRREI<br>CRIITKGTQTYSVLEGDPSENYSKYLLSLKEKEEDSSHTRAYGVCFVDTSLGK<br>FFIGQFSDDRHCSRFRTLVAHYPPVQVLFEKGNLSKETKTILKSSLSCSLQEG<br>LIPGSQFWDASKTLRTLLEEEYFREKLSDGIVMLPQVLKGMTSESDSIGLTPG<br>EKSELALSALGGCVFYLKKCLIDQELLSMANFEEYIPLDSDTVSKAYQRMVLD<br>AVTLNNLEIFLNGTNGSTEGTLLERVDTCHTPFGKRLLKQWLCAPLCNHYAIN<br>DRLDAIEDLMVVPDKISEVVELLKKLPDLERLLSKIHNVGSPLKSQNHPDSRAI<br>MYEETTYSKKKIIDFLSALEGFKVMCKIIGIMEEVADGFKSKILKQVISLQTKNP<br>EGRFPDLTVELNRWDTAFDHEKARKTGLITPKAGFSDYDQALADIRENEQSL<br>LEYLEKQRNRIGCRTIVYWGIGRNRYQLEIPENFTTRNLPEYELKSTKKGCKR<br>YWTKTIEKKLANLINAEERRDVSLKDCMRRLFYNFDKNYKDWQSAVECIAVL<br>DVLLCLANYSRGGDGPMCRPVILLPEDTPPFLELKGSRHPCGDDFIPNDILIG<br>CEEEEQKAYCVLVTGPNMGGKSTLMRQAGLLAVMAQMGCYVPAEVCRLTP<br>IDRVFTRLGSTFFVELSETASILMHATAHSLVLVDELGRGTATFDGTAIANAVV<br>KELAETIKCRTLFSTHYHSLVEDYSQNVAVRLGHMACMTFLYKFIKGACPKS<br>YGFNAARLANLPEEVIQKGHRKAREFEKMNQSLRLFRE |
| MSH6 Mut: 2<br><br>450E<br>451C<br>452H<br>453K | PTVWYHETLEWLKEEKRRDEHRRRPDHPDFDASTLYVPEDFLNSCTPGMR<br>KWWQIKSQNFDLVICYKVGKFYELYHMDALIGVSELGL**ECHK**GNWAHSGFP<br>EIAFGRYSDSLVQKGYKVARVEQTETPEMMEARCRKMAHISKYDRVVRREI<br>CRIITKGTQTYSVLEGDPSENYSKYLLSLKEKEEDSSHTRAYGVCFVDTSLGK<br>FFIGQFSDDRHCSRFRTLVAHYPPVQVLFEKGNLSKETKTILKSSLSCSLQEG<br>LIPGSQFWDASKTLRTLLEEEYFREKLSDGIVMLPQVLKGMTSESDSIGLTPG<br>EKSELALSALGGCVFYLKKCLIDQELLSMANFEEYIPLDSDTVSKAYQRMVLD<br>AVTLNNLEIFLNGTNGSTEGTLLERVDTCHTPFGKRLLKQWLCAPLCNHYAIN<br>DRLDAIEDLMVVPDKISEVVELLKKLPDLERLLSKIHNVGSPLKSQNHPDSRAI<br>MYEETTYSKKKIIDFLSALEGFKVMCKIIGIMEEVADGFKSKILKQVISLQTKNP<br>EGRFPDLTVELNRWDTAFDHEKARKTGLITPKAGFSDYDQALADIRENEQSL |

| | |
|---|---|
| | LEYLEKQRNRIGCRTIVYWGIGRNRYQLEIPENFTTRNLPEYELKSTKKGCKR<br>YWTKTIEKKLANLINAEERRDVSLKDCMRRLFYNFDKNYKDWQSAVECIAVL<br>DVLLCLANYSRGGDGPMCRPVILLPEDTPPFLELKGSRHPCGDDFIPNDILIG<br>CEEEEQKAYCVLVTGPNMGGKSTLMRQAGLLAVMAQMGCYVPAEVCRLTP<br>IDRVFTRLGSTFFVELSETASILMHATAHSLVLVDELGRGTATFDGTAIANAVV<br>KELAETIKCRTLFSTHYHSLVEDYSQNVAVRLGHMACMTFLYKFIKGACPKS<br>YGFNAARLANLPEEVIQKGHRKAREFEKMNQSLRLFRE |
| MSH6 Mut: 3<br><br>450C<br>451H<br>452K<br>453E | PTVWYHETLEWLKEEKRRDEHRRRPDHPDFDASTLYVPEDFLNSCTPGMR<br>KWWQIKSQNFDLVICYKVGKFYELYHMDALIGVSELGL**CHKE**GNWAHSGFP<br>EIAFGRYSDSLVQKGYKVARVEQTETPEMMEARCRKMAHISKYDRVVRREI<br>CRIITKGTQTYSVLEGDPSENYSKYLLSLKEKEEDSSHTRAYGVCFVDTSLGK<br>FFIGQFSDDRHCSRFRTLVAHYPPVQVLFEKGNLSKETKTILKSSLSCSLQEG<br>LIPGSQFWDASKTLRTLLEEEYFREKLSDGIVMLPQVLKGMTSESDSIGLTPG<br>EKSELALSALGGCVFYLKKCLIDQELLSMANFEEYIPLDSDTVSKAYQRMVLD<br>AVTLNNLEIFLNGTNGSTEGTLLERVDTCHTPFGKRLLKQWLCAPLCNHYAIN<br>DRLDAIEDLMVVPDKISEVVELLKKLPDLERLLSKIHNVGSPLKSQNHPDSRAI<br>MYEETTYSKKKIIDFLSALEGFKVMCKIIGIMEEVADGFKSKILKQVISLQTKNP<br>EGRFPDLTVELNRWDTAFDHEKARKTGLITPKAGFSDYDQALADIRENEQSL<br>LEYLEKQRNRIGCRTIVYWGIGRNRYQLEIPENFTTRNLPEYELKSTKKGCKR<br>YWTKTIEKKLANLINAEERRDVSLKDCMRRLFYNFDKNYKDWQSAVECIAVL<br>DVLLCLANYSRGGDGPMCRPVILLPEDTPPFLELKGSRHPCGDDFIPNDILIG<br>CEEEEQKAYCVLVTGPNMGGKSTLMRQAGLLAVMAQMGCYVPAEVCRLTP<br>IDRVFTRLGSTFFVELSETASILMHATAHSLVLVDELGRGTATFDGTAIANAVV<br>KELAETIKCRTLFSTHYHSLVEDYSQNVAVRLGHMACMTFLYKFIKGACPKS<br>YGFNAARLANLPEEVIQKGHRKAREFEKMNQSLRLFRE |
| MSH6 Mut: 4<br><br>450H<br>451K<br>452E<br>453C | PTVWYHETLEWLKEEKRRDEHRRRPDHPDFDASTLYVPEDFLNSCTPGMR<br>KWWQIKSQNFDLVICYKVGKFYELYHMDALIGVSELGL**HKEC**GNWAHSGFP<br>EIAFGRYSDSLVQKGYKVARVEQTETPEMMEARCRKMAHISKYDRVVRREI<br>CRIITKGTQTYSVLEGDPSENYSKYLLSLKEKEEDSSHTRAYGVCFVDTSLGK<br>FFIGQFSDDRHCSRFRTLVAHYPPVQVLFEKGNLSKETKTILKSSLSCSLQEG<br>LIPGSQFWDASKTLRTLLEEEYFREKLSDGIVMLPQVLKGMTSESDSIGLTPG<br>EKSELALSALGGCVFYLKKCLIDQELLSMANFEEYIPLDSDTVSKAYQRMVLD<br>AVTLNNLEIFLNGTNGSTEGTLLERVDTCHTPFGKRLLKQWLCAPLCNHYAIN<br>DRLDAIEDLMVVPDKISEVVELLKKLPDLERLLSKIHNVGSPLKSQNHPDSRAI<br>MYEETTYSKKKIIDFLSALEGFKVMCKIIGIMEEVADGFKSKILKQVISLQTKNP<br>EGRFPDLTVELNRWDTAFDHEKARKTGLITPKAGFSDYDQALADIRENEQSL<br>LEYLEKQRNRIGCRTIVYWGIGRNRYQLEIPENFTTRNLPEYELKSTKKGCKR<br>YWTKTIEKKLANLINAEERRDVSLKDCMRRLFYNFDKNYKDWQSAVECIAVL<br>DVLLCLANYSRGGDGPMCRPVILLPEDTPPFLELKGSRHPCGDDFIPNDILIG<br>CEEEEQKAYCVLVTGPNMGGKSTLMRQAGLLAVMAQMGCYVPAEVCRLTP<br>IDRVFTRLGSTFFVELSETASILMHATAHSLVLVDELGRGTATFDGTAIANAVV<br>KELAETIKCRTLFSTHYHSLVEDYSQNVAVRLGHMACMTFLYKFIKGACPKS<br>YGFNAARLANLPEEVIQKGHRKAREFEKMNQSLRLFRE |

**Table S3: Sequences of MSH6 (Blue are sites of mutagenesis, yellow is mutagenesis).**

| ID | Mutated Sequence | Pose | Interacting Residues | Binding Energy (kcal/mol) | Ligand Efficiency (kcal/mol) | Pose RMSD | Average RMSD |
|---|---|---|---|---|---|---|---|
| WT | ----- | 9 | M452<br>K453 | -6.6000 | -1.3200 | 77.1015 | 77.1037 |
| 1 | 450K<br>451E<br>452C<br>453H | 6 | H453 | -11.6400 | -2.3280 | 71.1618 | 71.1630 |
| 2 | 450E<br>451C<br>452H<br>453K | 6 | H452<br>K453<br>G454<br>H458 | -6.2800 | -1.2560 | 80.9530 | 71.0936 |
| 3 | 450C<br>451H<br>452K<br>453E | 1 | P362<br>W365<br>H451<br>E453<br>G454 | -6.9900 | -1.3980 | 70.0696 | 70.0913 |
| 4 | 450H<br>451K<br>452E<br>453C | 4 | H450 | -6.5000 | -1.3000 | 65.3302 | 65.3351 |

**Table S4. MSH6 Structure Prediction Scores.**

| MSH6 | Method | Template Structure Provided | Multiple Sequence Alignment | Score Type | Score | ptm |
|---|---|---|---|---|---|---|
| Wild-Type | Chai-1 | Yes | Yes | Aggregate Score | 0.186 | 0.931 |
| Mut: 1 | Chai-1 | Yes | Yes | Aggregate Score | 0.186 | 0.931 |
| Wild-Type | Boltz-2 | No | Yes | Confidence Score | 0.855 | 0.854 |
| Mut: 1 | Boltz-2 | No | Yes | Confidence Score | 0.856 | 0.851 |

**Table S5. Predicted Experimental Features of the Wild-Type and Mutant MSH6 Protein.**

| ID | pI | Instability Index | Half-Life Mammalian | Half-Life Yeast | Aliphatic Index | Grand Average hydropathicity |
|---|---|---|---|---|---|---|
| WT | 6.79 | 39.15 (stable) | >20 hours | >20 hours | 85.23 | -0.338 |
| Mut: 1 | 6.68 | 39.86 (stable) | >20 hours | >20 hours | 84.91 | -0.352 |
| Mut: 2 | 6.68 | 40.23 (unstable) | >20 hours | >20 hours | 84.91 | -0.352 |
| 3 | 6.68 | 39.84 (stable) | >20 hours | >20 hours | 84.91 | -0.352 |
| 4 | 6.68 | 39.97 (stable) | >20 hours | >20 hours | 84.91 | -0.352 |

*Expasy ProtParam calculated the pI and half-life of the wild-type and engineered MSH6 proteins.*

### Supplementary Figures

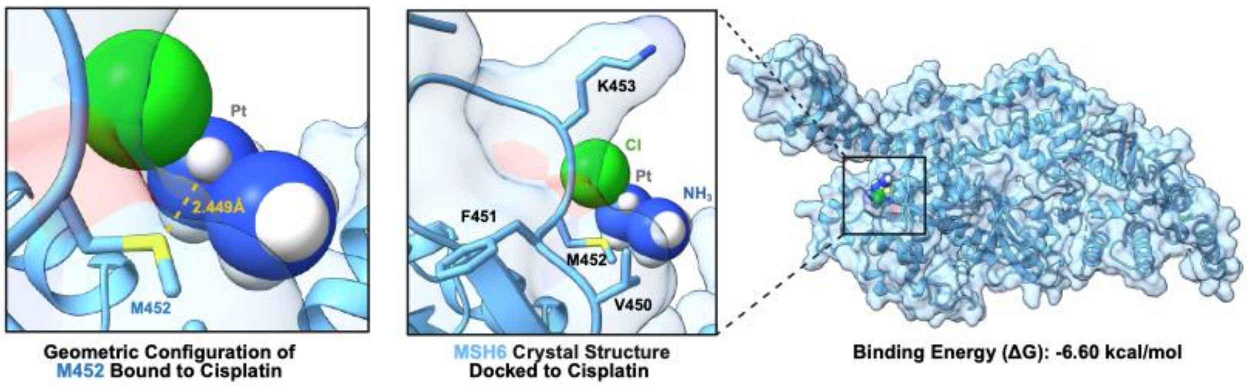


**Figure S1: Cisplatin Docked to the MSH6 Crystal Structure Using MetalDock.**

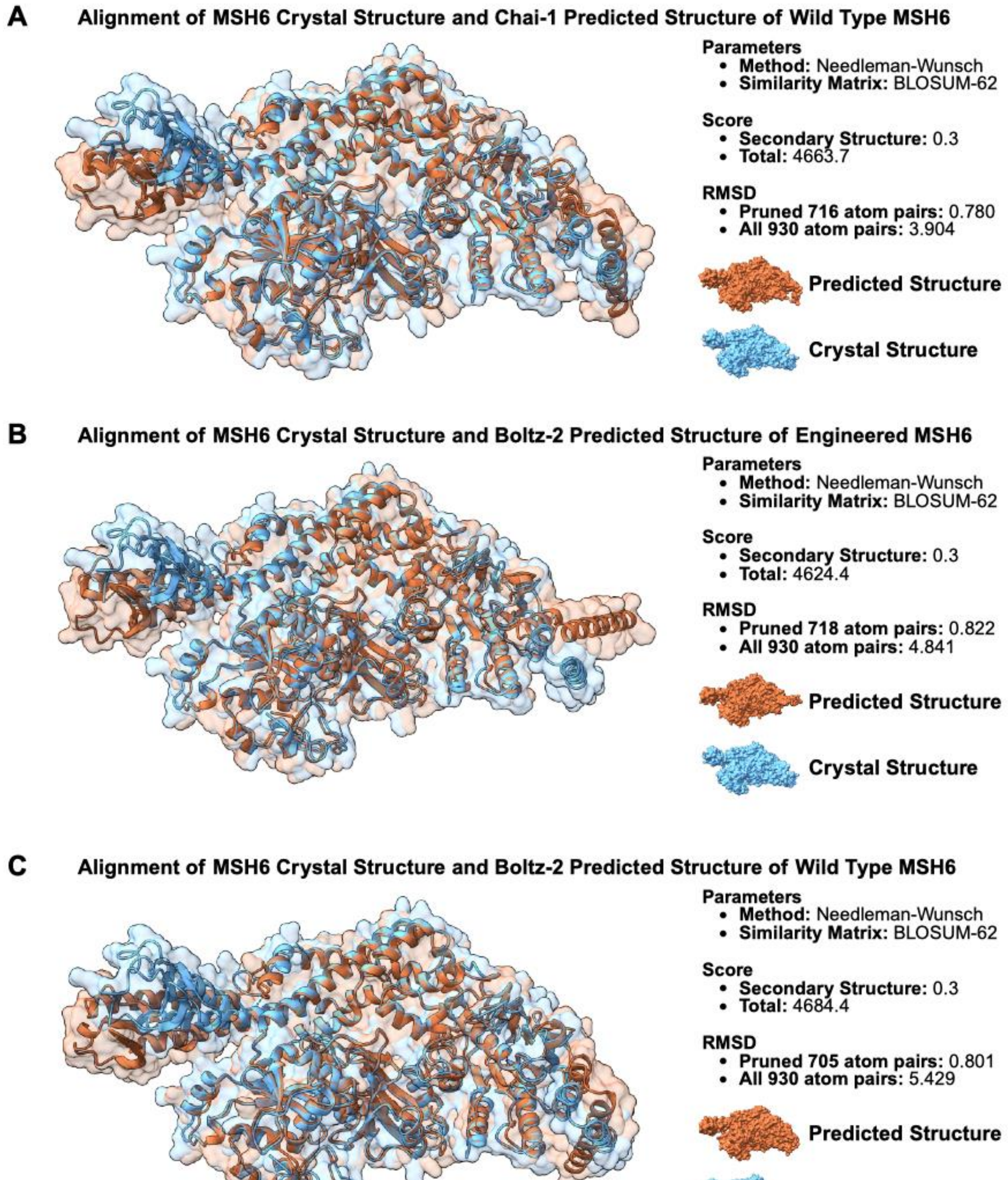


**Figure S2: Alignment of Predicted Structures of Wild Type and Engineered MSH6 to MSH6 Crystal Structure.**

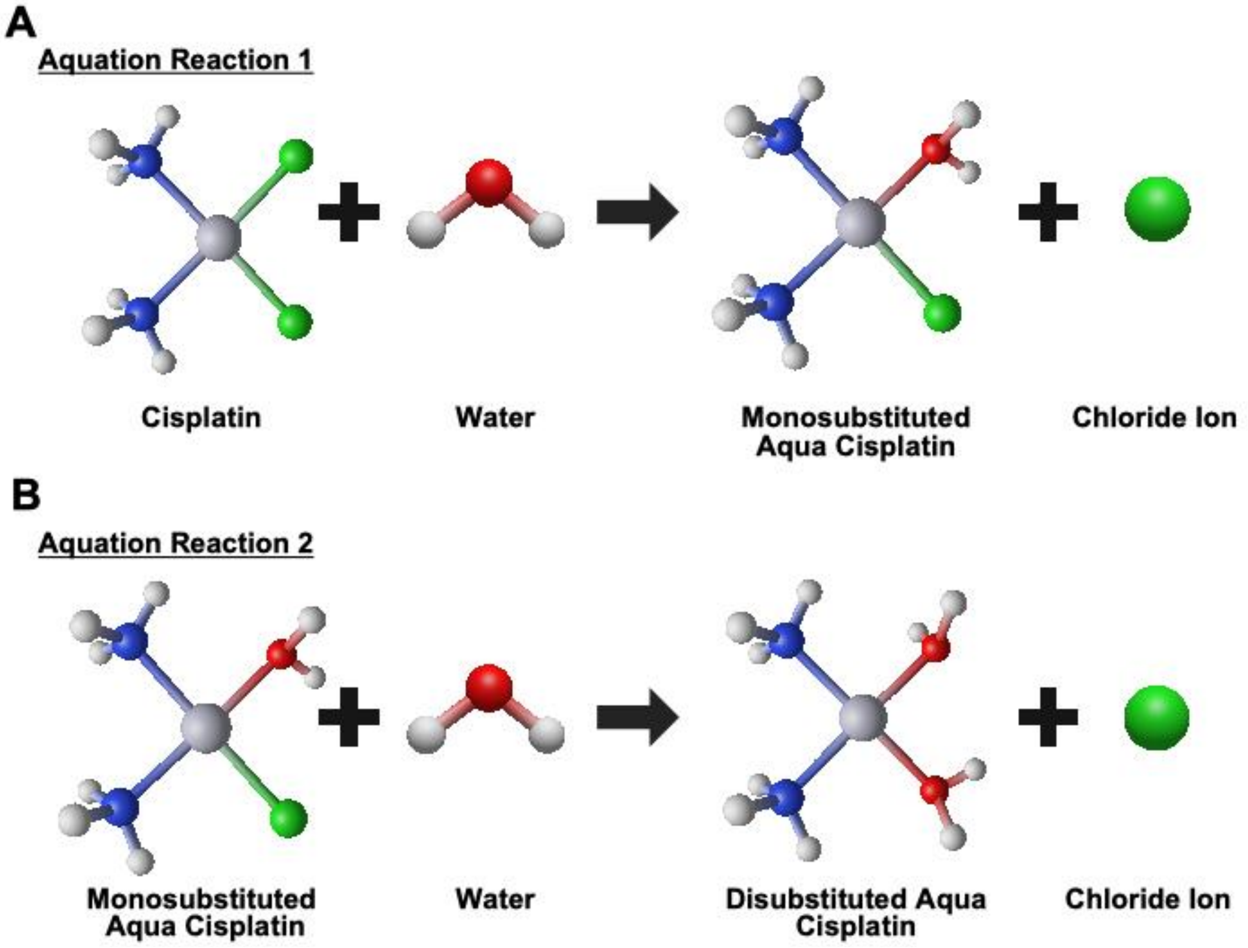


**Figure S3: Cisplatin Aquation Reactions Modeled with Geometrically Optimized Structures.**
